# Peyer’s Patch B cells undergo cell death via neutrophil-released toxic DNA following sterile tissue injury

**DOI:** 10.1101/2022.11.09.515763

**Authors:** Ali A Tuz, Alexander Beer, Markus Gallert, Dimitris Ttoouli, Susmita Ghosh, Sai P Sata, Andreas Kraus, Franziska Zwirnlein, Viola Kaygusuz, Vivian Lakovic, Altea Qefalia, Medina Antler, Sebastian Korste, Britta Kaltwasser, Hossam Abdelrahman, Ayan Mohamud-Yusuf, Chen Wang, Lars Haeusler, Smiths Lueong, Martin Stenzel, Benedikt Frank, Martin Köhrmann, Jens Siveke, Matthias Totzeck, Daniel Hoffmann, Anika Grüneboom, Nina Hagemann, Anja Hasenberg, Albert Sickmann, Jianxu Chen, Dirk M Hermann, Matthias Gunzer, Vikramjeet Singh

## Abstract

Lymphocyte contraction (LC) in central immune organs is a concomitant of sterile tissue injury, for example after stroke. Intestinal Peyer’s patches (PP) harbor large numbers of B cells, but how sterile tissue injury leads to LC in PP has not been explored. We observed rapid and macroscopically evident shrinkage of PP after stroke and myocardial infarction. Light-sheet fluorescence microscopy and flow cytometry revealed a strong reduction in the number of PP-resident B cells. Mechanistically, tissue injury triggered the activation of neutrophils that released B cell-toxic neutrophil extracellular traps (NETs) decorated with citrullinated histone-H3. Antibody-mediated or genetically induced neutrophil-loss, NETs-degradation or blockade of their generation completely reversed B cell loss and preserved the tissue architecture of PP. We also found NET-like elements in human post-stroke plasma. Hence, we propose that targeting NET-generation or -function counteracts post-injury B cell contraction in PP and thereby maintains immune homeostasis at mucosal barriers.

**In brief:** High numbers of B cells reside in the intestinal Peyer’s patches. Tuz et al. revealed that in response to sterile tissue injury, activated neutrophils release histone-decorated DNA into the circulation which induces B cell death. The loss of B cells results in the shrinkage of Peyer’s patches and reduced amounts of secretory IgA.

**Highlights:**

- Stroke and myocardial infarction induce the melting of Peyer’s patch
- Light-sheet microscopy and cytometry revealed B cell loss in Peyer’s patch
- Post-injury activated neutrophils release NETs and trigger B cell death
- Inhibition of NETs rescues B cell loss and degeneration of Peyer’s patch

## Introduction

Intestinal B cells are the major source of antibody-producing plasma cells and play a fundamental role in the protection of mucosal barriers. Among different gut-associated lymphoid tissues, Peyer’s patches (PP) are the most important site for the generation of B cells that later migrate to the lamina propria and secrete IgA (Lycke and Bemark, 2017). The generation of IgA-producing cells occurs in the germinal centers of PP and requires complex interactions between B cells, antigen-presenting cells, and helper T cells (Komban et al., 2019). The constant exposure of PP to commensal microflora or food antigens also induces tolerance in immune cells, thus inhibiting unwanted inflammation (Lin et al., 2021) and autoimmunity (Shirakashi et al., 2022). However, despite the important role of B cells in mucosal barrier defense and immune homeostasis, the impact of sterile tissue injury on intestinal B cells has not been extensively studied thus far.

Recent studies have shown that the loss of systemic T- and B cells after brain tissue injury can promote bacterial infections and deteriorate disease outcomes (Liesz et al., 2009; Roth et al., 2021; Singh et al., 2021). On the other hand, brain injury induces pro-inflammatory T cell subsets in PP that can migrate to the injured brain and increase neuroinflammation (Singh et al., 2016) which is accompanied by a reduction in the numbers of B and T cells in PP after cerebral ischemia (Schulte-Herbruggen et al., 2009). The current evidence indicates that, in contrast to inflammatory T cells, B cells positively influence functional recovery following brain injury. Consequently, preventing the loss of B cells along with maintaining defense against infections, might provide novel strategies for enhancing neurogenesis (Ortega et al., 2020). At present, such strategies have failed, mainly due to a lack of understanding of the underlying mechanisms of B cell loss from intestinal compartments after tissue injury.

The activation of sympathetic stress-signaling pathways after tissue injury can mediate the apoptosis of splenic B cells (McCulloch et al., 2017). However, clinical trials focusing on the blocking of stress-signaling pathways to inhibit immunosuppression and associated pneumonia in patients with brain injury have so far been less successful (Maier et al., 2018). To provide a novel explanation for massive brain-injury-associated systemic T cell loss, we recently demonstrated the contribution of increased circulating DNA (ciDNA) in inducing splenic T cell apoptosis via mechanisms involving inflammasome-dependent monocyte activation (Roth et al., 2021). However, whether ciDNA also influences the survival of B cells in PP has not been defined nor are the cellular source and molecular makeup of ciDNA clear.

In this study, we examined the consequences of brain- and myocardial injury on the tissue structure and cellular architecture of PP and revealed massive atrophy of intestinal PP and loss of local B cells. We showed that activated neutrophils released NETs as a major source of ciDNA which strongly triggered B cell death, PP architecture changes and eventually reduced amounts of secreted IgA.

## Results

### Tissue injury triggers contraction of Peyer’s patches via reducing B cell follicles volume

Tissue injury of vital organs is characterized by local necrosis that can impact the survival and functions of immune cells in distant lymphoid structures (Hoyer et al., 2019; Roth et al., 2021). To investigate whether tissue injury affects the structure of intestinal PP and functions of resident B cells, we employed a clinically relevant mouse model of ischemic stroke using transient middle cerebral artery occlusion (tMCAO). Following our previous studies, the induction of brain ischemia caused reproducible brain infarcts and behavioral deficits one and three days post-injury (Fig. S1, A-C). We identified that brain injury led to a marked reduction in the size of PP that was visible by the bare eye (Fig. 1A) and strongly reduced harvested PP numbers three days after brain injury (Fig. 1B). To further elucidate the extent of post-stroke PP disappearance, we adapted our 3D light sheet fluorescence microscopy (LSFM) protocols of the optically cleared murine intestinal tract (Schleier et al., 2020; Zundler et al., 2017) to perform a detailed volumetric analysis of PP (Fig. 1C and Fig. S1, D). The analysis confirmed a strong volume reduction (“melting”) of PP in brain-injury mice one and three days post-insult compared to sham controls (Fig. 1D). Previous studies have demonstrated that severe atrophy of spleen and thymus after brain injury was the result of local lymphocyte apoptosis (Offner et al., 2006). To analyze whether PP shrinkage was mediated via the loss of resident immune cells, we employed whole-mount immunostaining before LSFM analysis. Briefly, intestinal tissues with PP were stained with fluorescence-conjugated CD19 and CD3 antibodies and optically cleared before LSFM imaging (Fig. S1, D-F and video S1). The 3D reconstruction showed a strong shrinkage of PP follicular structures containing CD19^+^ B cells after brain injury (Fig. 1E). For a thorough and consistent analysis of large numbers of samples, we developed machine learning-based 3D volumetric image analysis (Fig. S1, G). Our data demonstrated that PP of brain-injury mice exhibited significantly smaller volumes of CD19^+^ B cell follicles in the jejunum and ileum compared to similar regions in sham controls (Fig. 1F and video S2). In contrast, brain injury caused a significant reduction of CD3^+^ T cell zone volumes only in the ileum (Fig. S2, A) with the general T cell zone volume in all PP being consistently much less than 50% of the B cell follicles volume, also under healthy conditions.

**Figure 1.**
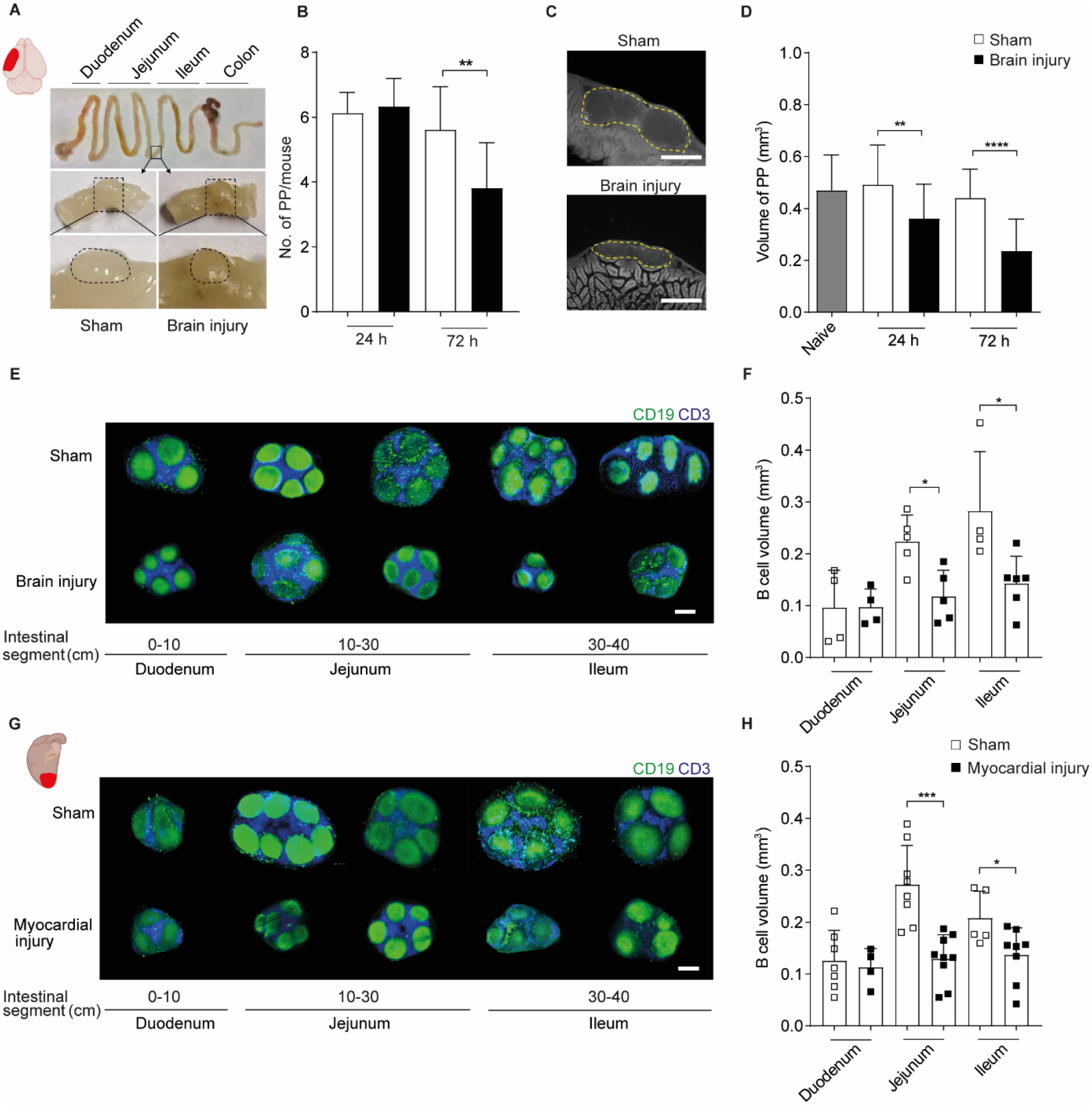
Sterile tissue injury induces shrinkage of intestinal Peyer’s patches (PP). **(A)** Macroscopic overview of the mouse gastrointestinal tract with the demarcation of PP one day after sham surgery or brain injury induced by tMCAO. **(B)** The total number of intestinal PP per mouse one and three days after sham or brain injury (n=8-13 mice per group). **(C)** Representative LSFM images of cleared intestinal PP showing their shrinkage one day after sham or brain injury. (**D)** Tissue volume analysis of PP after one and three days of sham or brain injury; (n=23 PP for sham and n=16 PP for brain injury, n=3-4 mice for one day and n=34 PP for sham and n=24 PP for brain injury, n=6 mice for three days). **(E)** 3D reconstruction LSFM images of CD19^+^ B cells and CD3^+^ T cells in PP from duodenum, jejunum and ileum after one day of brain injury or sham controls. **(F)** Deep learning-based automated analysis of B cell follicles volume in duodenum, jejunum and ileum after one day of brain injury or sham controls. (n=4-6 PP per intestinal segment). **(G)** 3D reconstruction images of CD19^+^ B cells and CD3^+^ T cells in PP derived from duodenum, jejunum and ileum one day after sham or myocardial injury. **(H)** Deep learning-based quantification of B cell follicles volume in duodenum, jejunum and ileum one day after sham or myocardial injury (n=4-9 PP per intestinal segment). Data are mean ± s.d., statistical analyses were performed by two-tailed Mann-Whitney test, *P<0.05, **P<0.01, ***P<0.001, ****P<0.0001. All data are combined from at least three independent experiments. PP=Peyer’s patches, Scale bar, 500 μm.

To test, whether these findings were specific to brain injury or rather a global response of PP B cells to large-scale tissue injury, we studied murine myocardial ischemia-reperfusion injury (Merz et al., 2019; Michel et al., 2021). Interestingly, the induction of myocardial injury also strongly reduced the size of PP follicular structures (Fig. 1G) and the volumes of CD19^+^ follicles in the jejunum and ileum after one day of surgery compared to sham controls (Fig. 1H). A similar reduction in CD3^+^ T cell zone volumes in PP was observed in the jejunum (Fig. S2, B). Collectively, these results showed a strong effect of sterile tissue injury on the rapid and massive loss of PP-resident lymphocytes and especially B cells.

### Tissue injury reduces B cells in PP and secretory amounts of IgA

To further quantify the influence of brain tissue injury on specific lymphocytes, we investigated their total numbers in PP using flow cytometry (Fig. S2, C). Our results showed that the majority of immune cells in PP were CD19^+^ B (78 ±10%) and CD3^+^ T (18 ±5%) cells with a small fraction of CD11b^+^ Ly6G^−^ monocytes (<2%) and Ly6G^+^ CD11b^+^ neutrophils (<1%) (Fig. S2, D). As expected, brain injury strongly reduced the total numbers of CD19^+^ B cells and CD3^+^ T cells in PP after one day (Fig. 2A). In keeping with our previous data (Roth et al., 2021; Singh et al., 2021), brain injury strongly decreased the numbers of B and T cells in spleen compared to controls (Fig. S2, E). Our flow cytometry analysis revealed that brain injury equally reduced the number of IgD^−^ germinal center (GC) and IgD^+^ pre-GC B cells in PP (Komban et al., 2019; Reboldi et al., 2016) (Fig. 2, B). In PP, B cells are primed by signals from the intestinal microflora and other immune cells to produce IgA- and IgG-secreting plasma cells (Clancy-Thompson et al., 2019; Reboldi et al., 2016). Interestingly, brain injury significantly decreased the number of IgA^+^ and IgA^−^ IgG1^−^ cells but did not influence the numbers of IgG1^+^ B cells (Fig. 2, C). Correspondingly, brain-injured mice showed significantly reduced amounts of secretory IgA in plasma and feces compared to sham controls (Fig. 2 D, E). However, the amounts of secretory IgG and IgM in plasma and feces remained unchanged.

**Figure 2.**
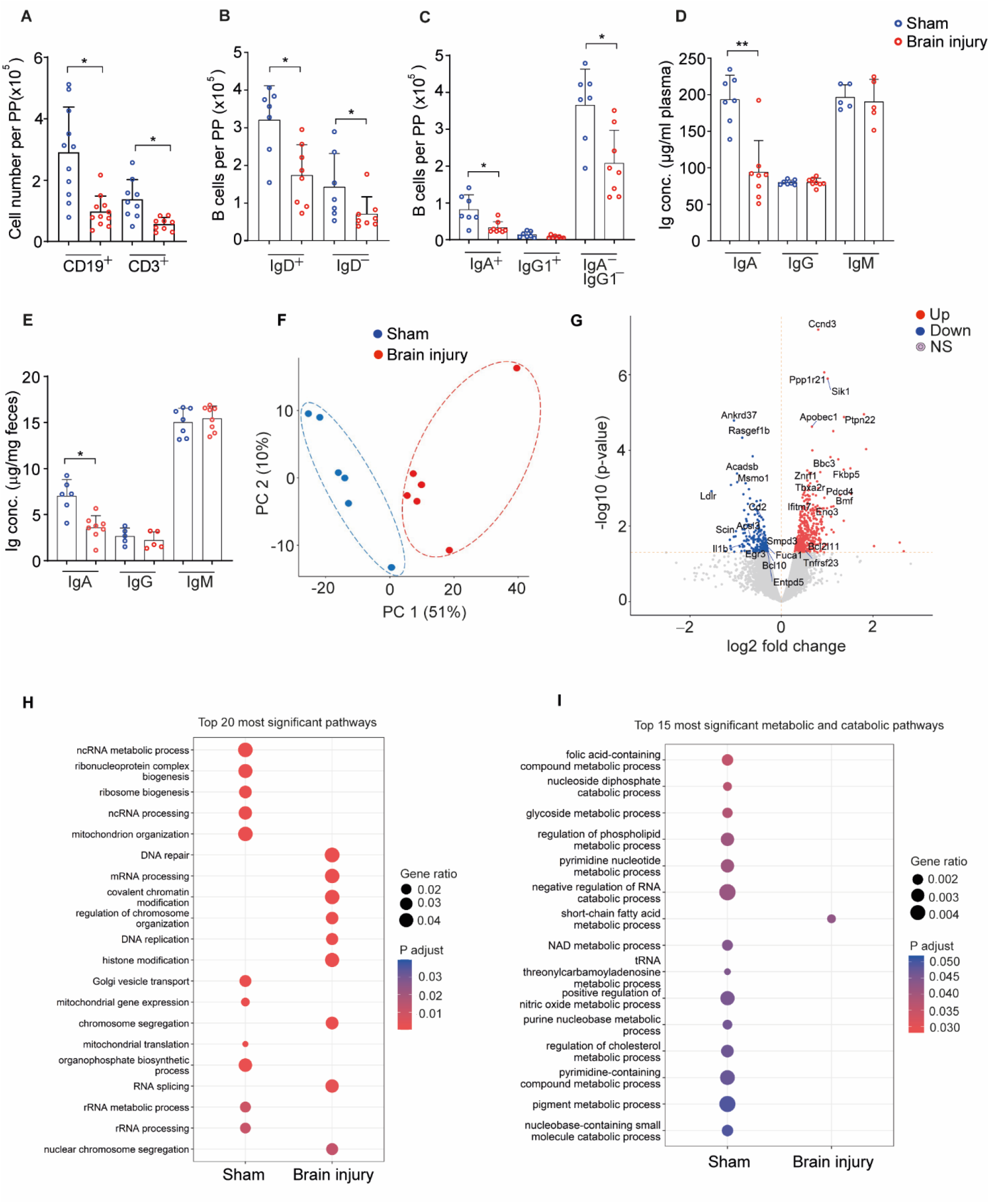
Brain tissue injury induces B cell loss in PP via activation of cell death pathways. **(A)** The flow cytometry-based quantification of the number of CD19^+^ B cells and CD3^+^ T cells in intestinal PP after one day of sham surgery or brain injury, (n=11-13 mice per group)**. (B)** The quantification of GC IgD^−^ and pre-GC IgD^+^ B cells in PP. **(C)** The quantification of IgA^+^, IgG1^+^ and IgA^−^IgG1^−^ B cells in PP. **(D)** The amount of plasma IgA, IgG and IgM in sham and brain–injured mice. (**E**) The amount of fecal IgA, IgG and IgM in sham–operated and brain-injured mice. **(F)** Principal component analysis (PCA) plot of RNA-seq data of CD19^+^ B cells in PP from mice exposed to sham surgery or brain injury (n=6 mice per group). **(G)** Volcano plot showing differentially expressed genes in PP B cells of sham-operated and brain-injured mice. Red dots indicate significantly upregulated genes and blue dots indicate significantly downregulated genes. **(H)** The gene set enrichment analysis (GSEA) shows enriched cell function pathways in sham-operated and brain-injured mice. **(I)** The GSEA shows enriched metabolic/catabolic pathways in PP B cells of sham-operated and brain-injured mice. Dot size indicates the calculated gene ratio and dot color indicates p–value representing the enrichment score as described in the methods. Data represent mean ± s.d., two-tailed Mann-Whitney test: *P<0.05, **P<0.01. All data is combined from at least three independent experiments, GC=germinal center.

### Peyer’s patches B cells undergo cell death after brain injury

We next aimed to investigate the molecular mechanism of B cell loss in PP after brain injury. Therefore, we purified PP B cells from mice 18 h after sham-operation or brain injury and analyzed transcriptomic changes by bulk RNA sequencing and bioinformatic analysis. Interestingly, the principal component analysis (PCA) showed that the transcriptomes of B cells clustered separately in brain-injured mice and sham controls (PC1, at 51%) (Fig. 2F). Moreover, 447 genes were upregulated and 198 genes were downregulated (absolute fold change ≥2 and adjusted p-value <0.05) in B cells following brain-injury compared to sham controls (Fig. 2G and table S1 and S2). As expected, cell death genes (Bmf, Pdcd4, Bbc3 and Znrf1) (Taylor et al., 2022; Woess et al., 2015) were enriched and cell function genes (Egr3, Ankrd37, Rasgef1b) and cell metabolic genes (Fuca-1, Smpd3, Acsl3, Ldlr and Entpd5) were reduced in B cells of brain-injured compared to sham control mice (Fig. 2G).

B cells from brain-injury mice expressed higher levels of the Myc target gene cyclin Ccnd3 and B cell lymphoma Bcl2, a key regulator of cell clonal expansion (Ramezani-Rad et al., 2020) and survival (Kurschat et al., 2021), respectively (Fig. 2G). In addition, brain injury enriched the expression of Sik1 and Ptpn22 in B cells which are involved in cell differentiation and B cell receptor signaling (Dai et al., 2013; Emily Robinson, 2018). These findings indicated that in parallel to the activation of pro-apoptotic genes, some B cell subsets may have upregulated survival genes after brain injury, potentially as compensatory mechanisms. Furthermore, we performed gene set enrichment analysis to identify cellular processes that were affected in B cells after brain injury. The differentially expressed genes enriched in B cells of brain-injured mice were related to chromatin and histone remodeling, DNA replication and repair (Fig. 2H). Notably, the pathways responsible for mitochondrial organization and function, ribosome biogenesis, and Golgi vesicle transport were decreased in B cells of brain-injured mice, suggesting severe metabolic disturbances in these cells. Further analysis showed a complete downregulation of pathways related to several cell metabolic and catabolic processes in B cells after brain injury compared to sham controls (Fig. 2I). Collectively, these data identified several key transcriptional changes in PP B cells after distant tissue injury and highlights severe immune disturbances at the intestinal mucosal tissues.

### Post-injury released circulating DNA prompts B cell loss in PP

While the loss of PP B cells after tissue injury was a robust finding, the underlying mechanisms remained unsolved. To elucidate this phenomenon, we focused on our earlier concept of soluble mediators released after tissue injury (Roth et al., 2021) and therefore analyzed their levels. The induction of brain injury in mice upregulated the levels of the stress hormone epinephrine (2 ng/ml vs 38 ng/ml), an indicator of sympathetic activation (Prass et al., 2003). Plasma epinephrine increase was observed 6 h after brain injury and remained high until the end of the study (24 h) (Fig. 3A). In line with our previous findings (Roth et al., 2021), we also found an early (6 h) increase in circulating DNA (ciDNA, 300 ng/ml vs 500 ng/ml) in brain-injured mice compared to controls (Fig. 3B). In contrast to 6 h, ciDNA amounts were decreased 24 h after brain injury but still significantly higher than in sham controls. Similarly, also myocardial injury increased the levels of ciDNA at 6 h which returned to sham levels at 24 h (Fig. S2, F). To test a potential causative role of adrenergic signaling and ciDNA for B cell loss in PP, we treated the mice with the β-receptor blocker propranolol or recombinant DNase-I immediately after brain injury. Interestingly, the degradation of ciDNA significantly inhibited the loss of PP B cells after brain injury while propranolol treatment was ineffective (Fig. 3C). Moreover, DNase-I treatment did not affect brain infarct volumes (Fig. 3D) at this early time point of 24 h post-insult, thus indicating a direct blockage of B cell loss with ciDNA acting as a soluble mediator released after brain injury.

**Figure 3.**
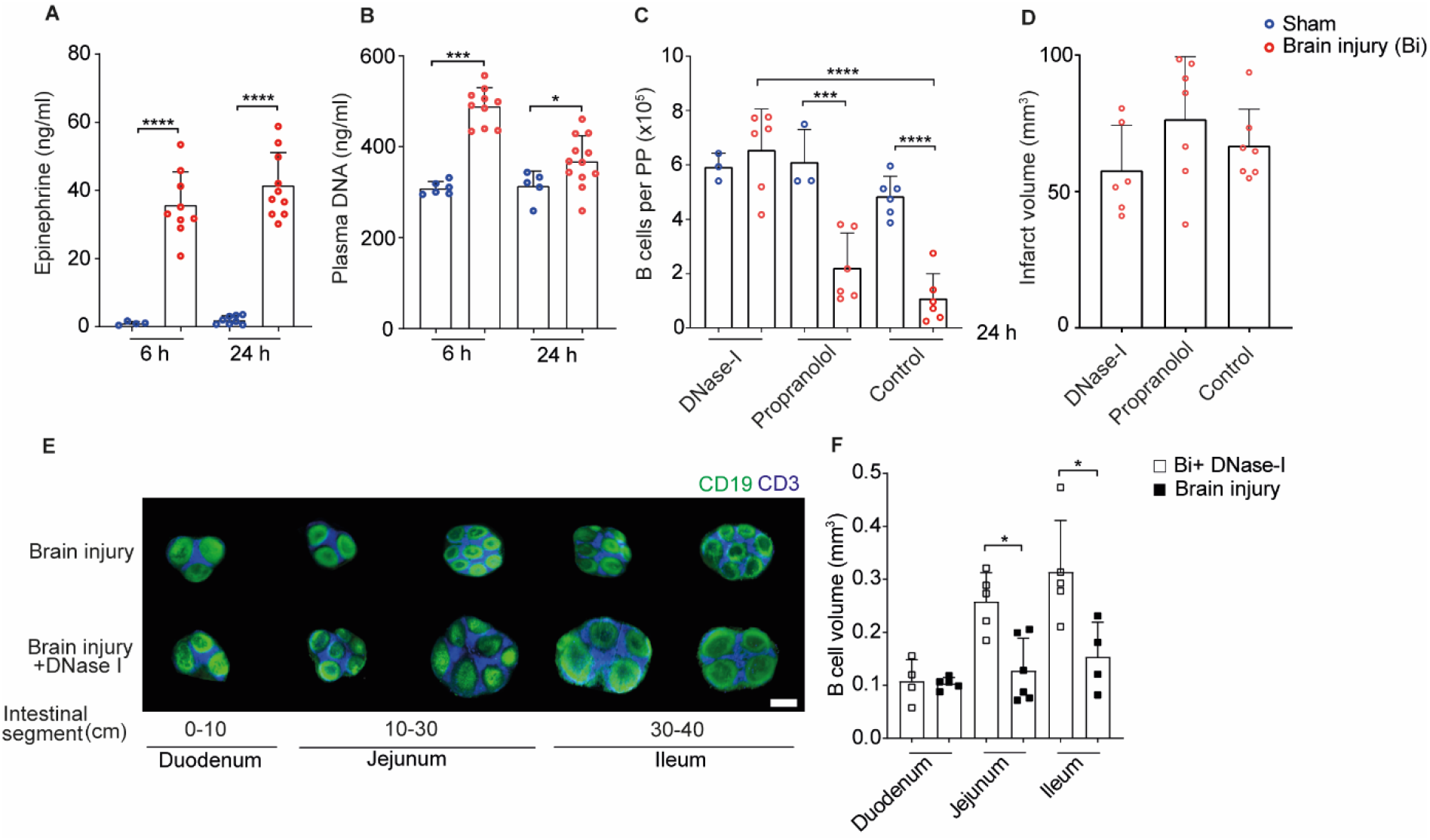
Brain injury-induced release of ciDNA promotes B cell loss in PP. **(A)** Quantification of plasma epinephrine and **(B)** DNA at 6 h and 24 h after sham surgery or brain injury (n=5-12 mice per group). **(C)** Numbers of B cells in PP 24 h after sham-operation or brain injury in DNase-I, propranolol and vehicle-treated mice. **(D)** Brain infarct volumes in DNase-I, propranolol treated and untreated mice at 24 h (n=6-7 mice per group). **(E)** 3D reconstruction images of CD19^+^ B cells and CD3^+^ T cells in PP after brain injury and DNase-I treatment. **(F)** The quantification of CD19^+^ B cell follicles volume in duodenum, jejunum and ileum 24 h after brain injury or brain injury + DNase-I treatment (n=4-6 PP per intestinal segment). Data are mean ±s.d., *p<0.05, ****P<0.0001, Shapiro-Wilk normality, and ordinary one-way ANOVA with Bonferroni’s multiple comparisons tests. PP= Peyer’s patches, Bi= Brain injury, Scale bar 500 μm.

Since ciDNA degradation with DNase-I prevented B cell death in PP, we investigated whether this treatment also preserved the mesoscopic structure of intestinal PP in brain-injured mice. LSFM analysis indeed confirmed that DNase-I treatment after brain injury completely preserved PP integrity (p<0.05) (Fig. 3E), and the volume of B cell compartments in the affected intestinal segments (Fig. 3F).

### Activated neutrophils produce circulating DNA as NETs after tissue injury

Next, we explored the potential sources of the additional amounts of ciDNA after brain injury. Activated neutrophils are the first intruders to injured inflammatory tissues and respond via the release of neutrophil extracellular traps (NETs) (Brinkmann et al., 2004).

To find out, whether circulating neutrophils in stroke mice were activated, we characterized their molecular makeup 6 h after sham surgery or brain injury using mass spectrometry (Fig. 4A). The comparative and label-free quantitative proteomic analysis yielded 1.757 proteins with ≥ 2 unique peptides and 1% FDR. A sample-wise comparison showed a Pearson correlation coefficient of ≥0.93, indicating good technical reproducibility (Fig. S3, A). Interestingly, the PCA showed a clear separation between circulating neutrophils of sham and brain-injured mice (Fig. 4B) based on the differential regulation of 89 proteins (*p*-value <0.05 and fold change ≥ 1.5). Of the 89 proteins, 32 were up- and 57 proteins were down-regulated in neutrophils of brain injury mice compared to sham controls (Fig. 4C and table S3, S4). Remarkably, proteins associated with neutrophil degranulation (Lamp1, S100a8, CD47) and neutrophil activation (Il1r2, Stat3) were upregulated in brain-injured mice (Fig. 4C). In addition, the increased abundance of apoptotic proteins (Aifm1, Casp3) and deficiency of cell function proteins (Pla2g7, Spta1, Sptb, Ipo5, Bsg, Gp1ba, Ptk2b) in brain-injured mice indicated the activation of cell-death pathways in neutrophils that have been observed in association with chromatinolysis (Tang et al., 2019) and NETosis (Sprenkeler et al., 2022; Vorobjeva and Chernyak, 2020).

**Figure 4.**
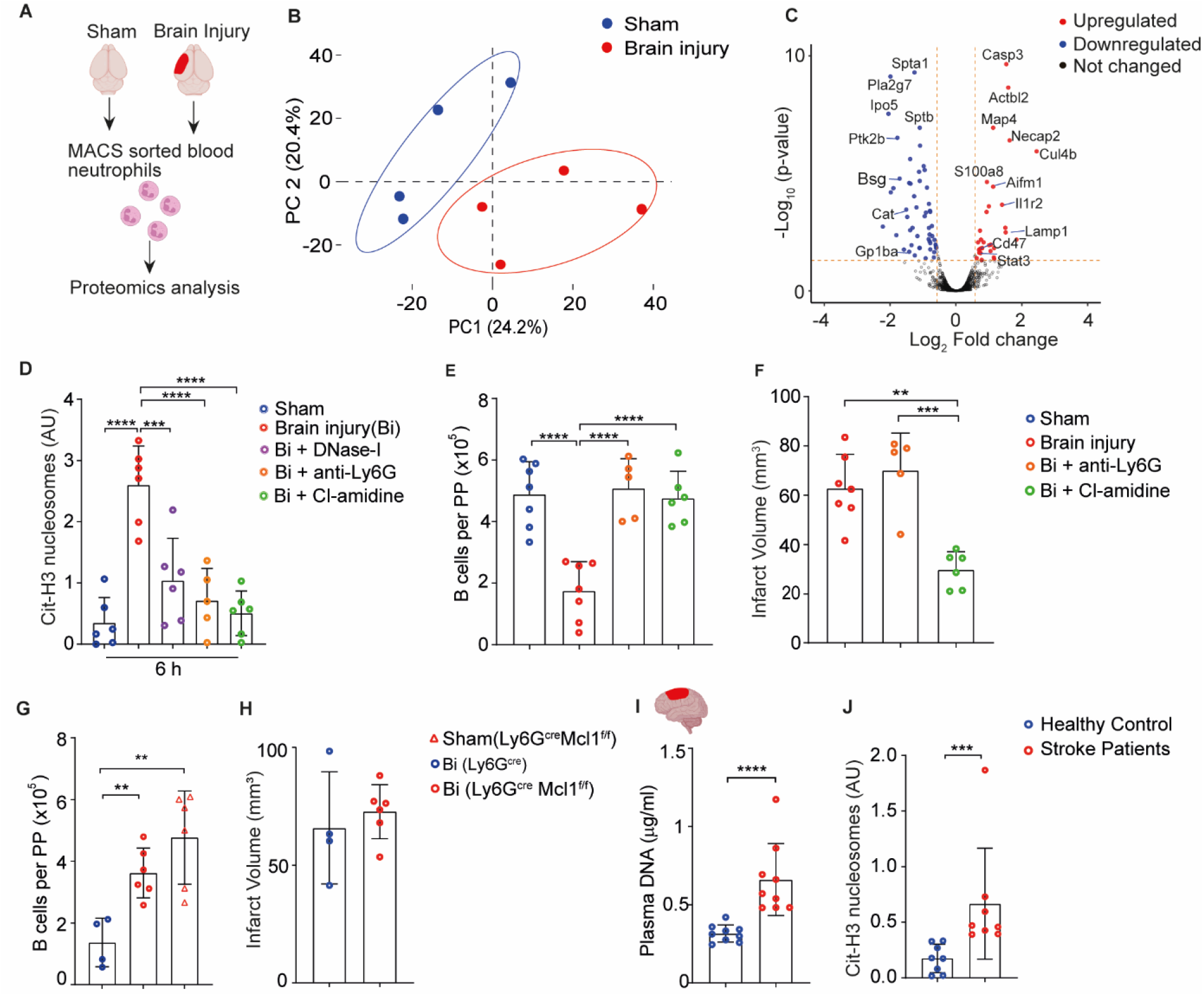
(A) Brain injury primes circulating neutrophils to release NETs in mice and stroke patients. Scheme of the experimental paradigm for neutrophil proteomics analysis. Blood neutrophils were isolated 6 h after sham surgery or brain injury to perform mass-spectrometry and proteomics analysis. **(B)** PCA of neutrophil proteomics after sham or brain injury (n=4 mice per group). **(C)** Volcano plot showing the analysis of differentially expressed proteins from blood neutrophils of sham-operated and brain injury mice. Red dots indicate significantly upregulated proteins and blue dots indicate significantly downregulated proteins. **(D)** Relative plasma levels of cit-H3 bound DNA after sham, brain injury, brain injury + DNase-I treatment or brain injury + anti-Ly6G antibody-treatment and brain injury + Cl-amidine. **(E)** Numbers of CD19^+^ B cells in intestinal PP in sham-operated, untreated brain injured and brain injured + anti-Ly6G antibody-treated and brain injury + Cl-amidine treated mice. **(F)** Brain infarct volumes in untreated brain injury, brain injury + anti-Ly6G treated and brain injury + Cl-amidine mice at 24 h (n=5-7 mice per group). **(G)** Numbers of CD19^+^ B cells in PP in sham-operated Ly6G^cre^Mcl1^f/f^ mice, brain-injured Ly6G^cre^ and Ly6G^cre^ Mcl1^f/f^ mice. **(H)** Brain infarct volumes in brain-injured Ly6G^cre^ and Ly6G^cre^Mcl1^f/f^ mice at 24 h (n=4-6 mice per group). **(I)** Quantification of plasma DNA in ischemic stroke patients and healthy controls. **(J)** Relative plasma levels of cit-H3 bound DNA in stroke patients within three days of admission and healthy controls (n=8-9 per group). D, E, F: Data represent mean ±s.d., ***p<0.001, ****P<0.0001, Shapiro-Wilk normality, and ordinary one-way ANOVA with Bonferroni’s multiple comparisons tests. G, I, J: Data are mean ±s.d., **p<0.01, ***p<0.001, ****P<0.0001, two-tailed Mann-Whitney test. PP=Peyer’s patches, Bi= Brain injury.

To determine whether NETs were indeed released into the circulation after brain injury, we measured the plasma content of citrullinated histone H3 (cit-H3), a hallmark of NET-DNA (Tsourouktsoglou et al., 2020). Indeed, 6 h after brain injury, we observed increased levels of citH3-containing nucleosomes in blood (Fig. 4D). Interestingly, treatment of mice with DNase-I immediately after brain injury reduced citH3-containing nucleosome levels (Fig. 4D). To further validate the contribution of neutrophils to citH3 release and B cell loss in PP, we applied our established dual antibody-mediated depletion of circulating neutrophils (Fig. S3, B) (Boivin et al., 2020). Indeed, neutrophil removal before the induction of brain injury substantially reduced citH3-containing plasma nucleosomes (Fig. 4D) and total DNA in plasma (Fig. S3, C) and also completely inhibited the loss of B cells in PP (Fig. 4E), without affecting the brain infarct volumes (Fig. 4F). The enzyme PAD4 (peptidyl arginine deiminase 4) has been shown to participate in the formation of NETs and inhibition of PAD blocks NETosis (Li et al., 2010). Treatment of mice with the PAD4 inhibitor Cl-amidine immediately after induction of brain injury significantly reduced plasma cit-H3 amounts at 6 hours (Fig. 4D) and prevented the loss of B cells in PP at 24 h (Fig. 4E). However, the administration of Cl-amidine after brain ischemia also reduced brain infarcts (Fig. 4F), showing additional neuroprotective effects of NET inhibition.

Acutely depleting millions of neutrophils by antibody injection might have uncontrolled secondary effects on immune homeostasis. Hence, we generated mice, where neutrophils were absent due to the genetic removal of the critical survival factor Mcl (Silvestre-Roig et al., 2019a). Confirming our antibody-depletion experiments, the genetically induced loss of neutrophils in Ly6g^cre^Mcl^f/f^ mice (Fig. S3, D), showed a higher number of PP B cells compared to neutrophil-sufficient Ly6g^cre^ mice 24 h after brain injury (Fig. 4G) without changing the infarct volumes (Fig. 4H). Moreover, neutrophil-deficient Ly6g^cre^Mcl^f/f^ mice also showed an increased number of splenic T cells compared to brain-injured Ly6G^cre^ controls, thus suggesting the contribution of neutrophils also in increased ciDNA generation and T cell apoptosis-induction as observed in our previous study (Fig. S3 E) (Roth et al., 2021). Since we did only find small numbers of Ly6G^+^ neutrophils or NET-forming citH3^+^ neutrophils directly within the PP of brain-injured mice (Fig. S3 F, G), we assume that acute NET production by circulating neutrophils in response to sterile injury is the trigger that leads to PP B cells.

To evaluate the relevance of our experimental findings for the clinical reality, we finally analyzed circulating NETs in the plasma of 10 human ischemic stroke patients that were collected within 3 d of stroke-unit admission. The blood samples from 9 equally aged healthy individuals were collected and processed with identical protocols. The plasma of stroke patients contained significantly higher levels of total DNA compared to healthy individuals (Fig. 4I) as shown previously (Grosse et al., 2022). Strikingly, however, also the levels of citH3-containing nucleosomes were significantly increased in the plasma of stroke patients compared to healthy individuals (Fig. 4J), thus suggesting NET-release in human patients following acute stroke.

## Discussion

The intestine undergoes significant morphological and immune alterations in response to brain tissue injury, with intestinal microbiota and motility disturbances (Singh et al., 2016), reduced mucus secretion (Houlden et al., 2016) and induction of pro-inflammatory T cell subsets (Benakis et al., 2016). Our findings that stroke and myocardial infarction trigger loss of B cells in intestinal Peyer’s patches (PP) expose an unrecognized link between tissue injury to intestinal immune dysbalance. The loss of B cells also revealed massive tissue shrinkage and later disappearance of PP after tissue injury. Previous studies have demonstrated massive atrophy of systemic lymphoid tissues such as spleen and thymus after brain tissue injury which is mediated via apoptosis of local immune cells (Liesz et al., 2009; Offner et al., 2006). B cells in PP are organized in follicular structures and generate large numbers of precursors for antibody-secreting plasma cells, and our data showed that B cell loss after brain tissue injury instigated reductions in the numbers of IgA-producing B cells and, as a result, secretory levels of IgA antibodies. We and others have provided evidence for intestinal T-cell involvement in neuroinflammation (Benakis et al., 2016; Singh et al., 2016) and systemic B-cell impact on neuroregeneration after brain injury (Ortega et al., 2020). B cells protect mucosal barriers and also have the capacity to synthesize neurotrophins to support neuronal survival (Ren et al., 2011). These effector functions of B cells are dependent on their survival, maintenance and differentiation and are regulated transcriptionally (Laidlaw and Cyster, 2021). We performed bulk RNA sequencing and provide evidence for severe alterations in the transcriptomics of B cells in PP of brain-injured mice compared to controls. Our analysis revealed changes in pathways related to cellular energy generation, DNA replication and several metabolic processes in B cells of brain-injured mice. Of note, various apoptotic, survival and proliferation pathway genes were regulated in B cells of brain-injured mice which indicates the activation and inhibition of multiple molecular pathways at the same time (Kurschat et al., 2021; Ramezani-Rad et al., 2020). The precise information on the modulation of different transcriptional pathways in a specific subset of B cells is not clear and will require single-cell RNA analysis. However, our single-cell flow cytometry data reveal the significant effect of brain injury on IgA and IgA^−^IgG1^−^ B cell subsets in PP.

Mechanistically, the massive loss of immune cells has been demonstrated to be partly dependent on the over-activation of sympathetic signaling pathways, which is indicated by the increased levels of systemic catecholamines (Prass et al., 2003). However, the clinical studies blocking this pathway using beta-blockers have not prevented immunosuppression nor improved stroke outcomes (Westendorp et al., 2016). In this regard, we have recently demonstrated the major contribution of circulating DNA (ciDNA) as a soluble mediator deriving apoptosis of splenic T cells via inflammasome-dependent macrophage activation (Roth et al., 2021). With the findings presented here the increased levels of ciDNA can also be linked to the loss of B cells in PP after sterile tissue injury. Our data revealed upregulated levels of ciDNA and epinephrine early after the induction of brain- and myocardial injury in mice. The finding that enzymatic degradation of ciDNA but not the blockage of beta-adrenergic signaling prevented loss of B cells and maintained the structural integrity of PP further highlights the key role of post-injury released NETs on intestinal immunosuppression and does not support a substantial effect of sympathetic activation on this process (Prass et al., 2003).

Soluble mediators released after sterile tissue injury have been shown to activate neutrophils that then induce the formation of toxic NETs (Denorme et al., 2022). NETs consist of DNA, histones, and granular proteins and increase collateral inflammation and cell death. Recently, we showed that NETs directly induce smooth muscle cell death in arterial inflammation via attached histone-H4 proteins (Silvestre-Roig et al., 2019b). Our proteomics results reveal neutrophil activation as a consequence of tissue injury as early as six hours after ischemic insult. Brain injury increases the abundance of neutrophil activation-associated proteins such as S100A8, Lamp1 and Casp3 that might facilitate NET formation (Sprenkeler et al., 2022) as well as Aifm1, which is associated with chromatinolysis (Tang et al., 2019).

Previous findings have shown that the extent of brain injury after cerebral ischemia is decisive for the degree of lymphoid immune cell loss (Liesz et al., 2009). In contrast, our findings on brain infarct size did not show a correlation of brain-tissue protection with NETs degradation or neutrophil depletion, at least in the first 24 h post-stroke. Moreover, since neutrophil deficiency also prevented the loss of splenic T cells after brain injury, this study now provides a likely explanation for the previously unidentified source of ciDNA as seen in our previous study (Roth et al., 2021). In recent years, DNase therapy has been largely proposed for the treatment of COVID-19 (Fisher et al., 2021) and psoriasis patients (Shao et al., 2019). Our findings that NETs are also present in the plasma of stroke patients, suggest clinical trials to test the efficacy of early DNase therapy to reduce immunosuppression and thereby improve disease outcomes. This would require broadening the focus of clinical stroke trials from just neuroprotective to immunoprotective effects, which might also have substantial positive effects for patients.

In summary, our results show that sterile tissue injuries lead to immediate degeneration (“melting”) of PP via the induction of NET release and the ensuing death of resident follicular B cells in mice. This appears to be a global response of intestinal B cells toward ciDNA released as an inflammatory soluble mediator in response to sterile tissue necrosis. However, a mechanistic understanding of how circulating NETs access PP and induce B cell apoptosis following tissue injury is not described by this study. Based on the presence of citH3 histones, the most likely source of post-infarction ciDNA are NETs (Brinkmann et al., 2004). Although previously speculated (Grosse et al., 2022), this has not been experimentally proven until now. Our findings that depletion of neutrophils fully restores the integrity of PP makes it less likely that ciDNA released from the decaying brain is responsible for immune cell loss. Previously, NETs have mainly been associated with impairment of post-stroke brain recovery (Kang et al., 2020). Here, we show significant early effects on immune cell survival and function that open attractive treatment options by targeting NET-release or -function, for example as emergency therapy in patients with stroke or myocardial infarction.

### Limitations of the study

In our study, the depletion of neutrophils or degradation of NETs did not affect the primary brain tissue injury in the period of 24 h, in which PP follicular B cell loss was observed. Future studies will have to address the long-term impact of NETs inhibition on brain tissue protection along with intestinal immune homeostasis. Furthermore, RNA-sequencing analysis was only performed on a pool of B cells, the results may derive from different subsets with heterogeneous transcriptional responses. However, in general, results are well in line with the apoptotic loss of follicular B cells observed by flow cytometry and microscopy.

## Acknowledgments

This research work was funded by the Deutsche Forschungsgemeinschaft (DFG), research grant SI 2650/1-1 to VS, GU 769/10-1 to MG, HE 3173/11-1, 3173/12-1, 3173/13-1 and 3173/15-1 to DMH and the CRC TRR332 (project C6) to MG and DMH. The work of SS and JC was supported by the Bundesministerium für Bildung und Forschung, BMBF, Ref. 161L0272. The microscopy and flow cytometry works in this manuscript were performed at the Imaging Center Essen (IMCES) a service core facility of the faculty of medicine of the University Duisburg-Essen, Germany. We thank Ms. Alexandra Brenzel and Dr. Anthony Squire at IMCES for their technical support. All authors have read and approved the manuscript, and the manuscript has not been accepted or published elsewhere.

## Supplementary data

**Figure S1.**
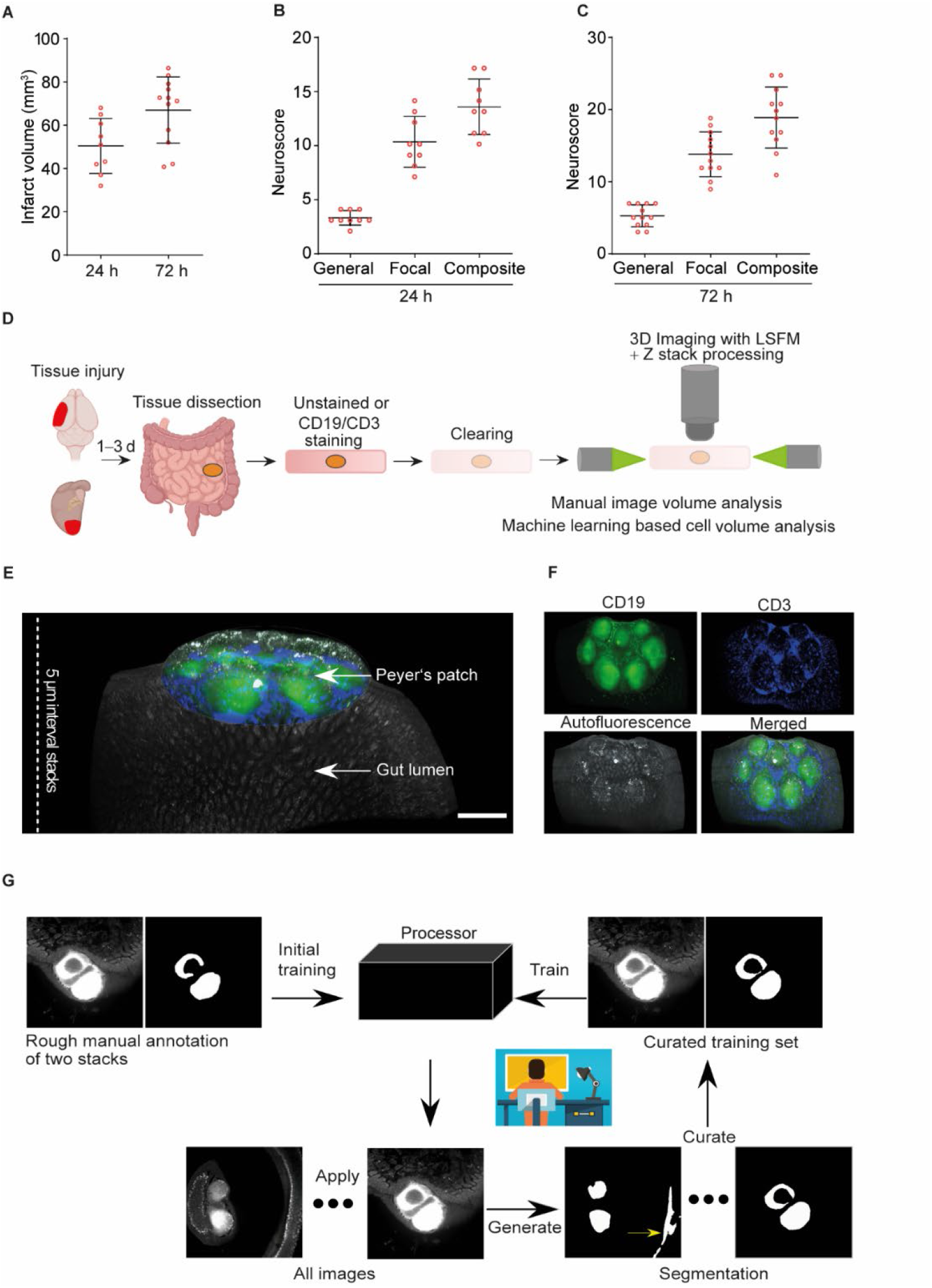
Brain injury induces sensorimotor deficits and its impact on PP B cell follicles volume is analyzed using LSFM. **(A)** Quantification of brain infarct volume after one and three days of brain injury. **(B)** Neurological deficits are measured on one day and **(C)** three days after brain injury (n=9-12 mice per group, data represented as mean ± s.d.). **(D)** Illustration of the intestinal tissue preparation for unstained-volume analysis of whole PP or whole-mount staining of B and T cells in PP, followed by tissue clearing and LSFM-based 3D volume analysis. **(E)** Fluorescence images after 3D reconstruction of stained PP showing the position of PP in the small intestine that were whole-mount stained with CD19 (green) and CD3 (blue) fluorescence antibodies before LSFM. **(F)** Fluorescence single-channel images after 3D reconstruction of stained PP showing CD19^+^ B cells (green), CD3^+^ T cells (blue), and autofluorescence signal (grey). **(G)** Overview of the human-in-the-loop segmentation workflow for automated analysis of B cell follicles volume and T cell zone volume. Scale bar, 500 μm.

**Figure S2.**
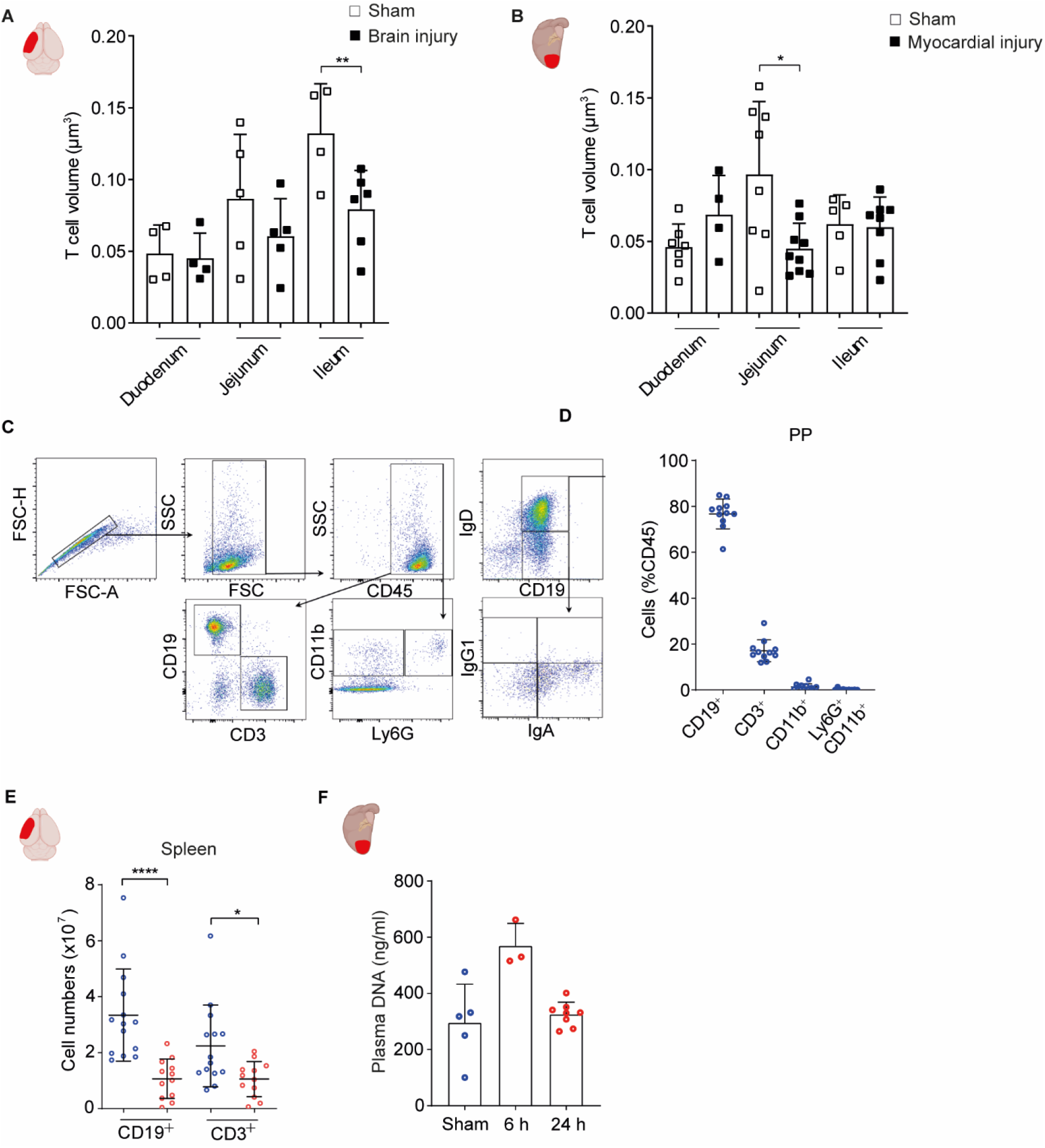
Tissue injury induces loss of B and T cells in PP and spleen. **(A)** The quantification of T cell zone volume in different intestinal segments after one day of brain injury or sham controls (n=4-6 PP per intestinal segment). **(B)** The quantification of T cell zone volume in different intestinal segments after one day of myocardial injury or sham controls (n=4-6 PP per intestinal segment). **(C)** Representative gating strategy for the flow cytometry analysis. **(D)** The frequency of CD19^+^, CD3^+^, CD11b^+^ and Ly6G^+^ CD11b^+^ cells in PP of sham mice. mice. **(E)** The quantification of CD19^+^ and CD3^+^ cells in spleen after 24 h of sham-operation or brain injury (n=11-13 mice per group). **(F)** The quantification of plasma DNA after 6 h of sham-operation or myocardial injury. Data represented as mean ± s.d., statistical analyses were performed by two-tailed Mann-Whitney test, *P<0.05, **P<0.01, ****P<0.01. All data are combined from at least three independent experiments.

**Figure S3.**
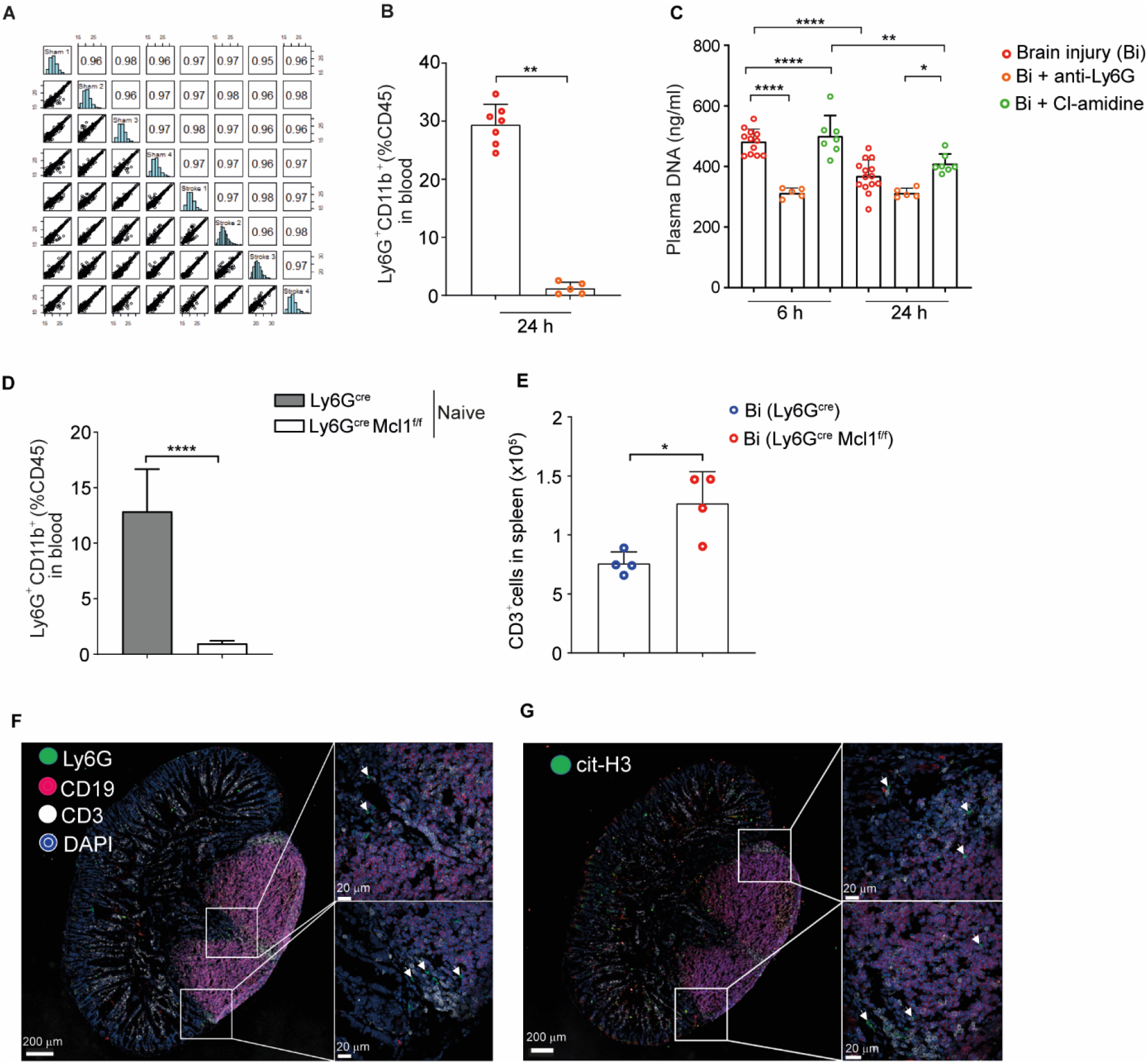
Neutrophil-released NETs contribute to increased plasma DNA after tissue injury. **(A)** A sample-wise comparison showed a Pearson correlation coefficient for all analyzed samples for mass spectrometry analysis. **(B)** The percentage of Ly6G^+^ leucocytes in blood in untreated and anti-Ly6G antibody-injected mice at 24 h (n=5-7 mice per group). **(C)** The quantification of plasma DNA at 6 h and 24 h after in untreated and anti-Ly6G antibody-injected and Cl-amidine treated mice (n=5-14 mice per group). **(D)** Percentages of blood Ly6G^+^ CD11b^+^ neutrophils in naïve Ly6G^cre^ and Ly6G^cre^Mcl1^f/f^ mice (n=11-15 mice per group). **(E)** The numbers of CD3^+^ cells in spleens of brain-injured Ly6G^cre^ and Ly6G^cre^Mcl1^f/f^ mice. **(F)** The representative fluorescence images of Ly6G and **(G)** cit-H3 stained PP after six hours of brain injury. Data represented as mean ±s.d., statistical analyses were performed by two-tailed Mann-Whitney test, *P<0.05, **P<0.01, ****P<0.0001. Bi=Brain injury.

