## Supplementary material for "Peyer’s Patch B cells undergo cell death via neutrophil-released toxic DNA following sterile tissue injury": Materials and Methods

### Mice

All animal experiments were performed following ethical guidelines and were approved by the local authorities (G1713/18; G1719/19; G1650/17) of the Landesamt für Natur, Umwelt und Verbraucherschutz Nordrhein-Westfalen, Recklinghausen, Germany). Male C57BL/6JHsd wild-type male mice were received from Envigo, Netherlands. For myocardial infarction experiments, male C57BL/6JRj wild-type male mice were obtained from Janvier Labs, France. Mice were randomly divided into two groups that underwent sham surgery or ischemic brain or myocardial injury. All experiments were performed and reported according to ARRIVE guidelines (Kilkenny et al., 2010).

### Mouse model of brain injury

Brain injury was induced by transient middle cerebral artery occlusion (tMCAO) in C57BL/6JHsd mice (aged 8-10 weeks) anesthetized with 1% isoflurane in 100% oxygen. Mice were injected with the analgesic buprenorphine (0.1 mg/kg body weight, s.c.) 30 min before the surgery. An eye ointment (Bepanthen) was applied to avoid any harm to mouse eyes during the surgical procedures. A small incision was made between the ear and the eye to expose the temporal bone and a laser Doppler flow probe was attached to the skull above the core of the middle cerebral artery (MCA) territory. Mice were then placed in a supine position on a feedback-controlled heat pad and the midline neck region was exposed with a small incision. The common carotid artery (CCA) and left external carotid arteries were identified and ligated. A 2 mm silicon-coated filament (Cat. 702234PK5Re; Doccol) was inserted into the internal carotid artery to occlude the MCA. Brain ischemia was validated by a stable reduction of blood flow to ≤ 20% of baseline that was observed on the laser Doppler flow device. After 60 min of occlusion, the filament was removed for the reestablishment of blood flow. Mice were then injected with anti-inflammatory drug carprofen (4-5 mg/kg body weight, s.c.) and wounds were carefully sutured and mice returned to their cages with free access to food and water. Sham-operated mice underwent the same surgical protocol except the filament was inserted into the CCA and immediately removed. For experiments with three days of survival, mice were daily injected with carprofen (4-5 mg/kg body weight, s.c.) and 5% glucose. Post-injury sensorimotor behavioral deficits were evaluated by the Clark neuroscore (Table S5). The exclusion criteria for experimental mice were as follows: inadequate ischemia (reduction of blood flow does not reach the ≤ 20% of baseline threshold), weight loss >20% of baseline body weight during the study, and spontaneous animal death.

**Mouse model of myocardial injury**

Male C57BL/6JRJ mice (aged 9-15 weeks) were subjected to a myocardial ischemia-reperfusion injury as previously described (Totzeck et al., 2016). Briefly, mice were anesthetized i.p. with ketamine (100 mg/kg) and xylazine (10 mg/kg). Mice were then orally intubated and ventilated throughout the operation procedure, with 0.8 l/min air and 0.2 l/min O2 at a tidal volume of 250 µl/stroke and a breathing frequency of 140 strokes/min. A supplementation of 2% isoflurane to ventilated air was used to maintain anesthesia. The chest was opened through a left lateral thoracotomy and the left coronary artery (LCA) was ligated. After 45 min of ischemia, the affected myocardium was reperfused. After 24 h, mice were sacrificed by isoflurane anesthesia and then cervical dislocation and blood samples were taken from the inferior vena cava. Mice were then perfused with 0.1 M phosphate buffer saline (PBS) supplemented with heparin (20 IE/ml) and different organs were collected in cold PBS and placed on ice for further processing.

**Quantification of brain infarct volumes**

For the calculation of brain infarct volumes, brains were carefully removed and quickly frozen on dry ice. Afterward, 20 µm thick cryosections were cut at 500 µm intervals and stained with cresyl violet (Cat. C5042, Sigma). All sections were scanned using 600 dpi and analyzed using the ImageJ software (NIH). Using a scale of 23.62 pixels/mm, the area of non-stained infarct tissue was measured and integrated into the total brain. An edema correction for brain infarct volume was performed using the following formula: (ischemic area) =(direct lesion volume) – [(ipsilateral hemisphere) – (contralateral hemisphere)]. The lesion volume per hemisphere was presented as $mm^3$.

**Single-cell preparations from lymphoid organs and flow cytometry analysis**

Mice were deeply anesthetized with Ketamine/Xylazine (100 mg/kg/10 mg/kg, i.p.) and blood was collected via cardiac puncture and added to EDTA-containing tubes. Mice were then perfused with PBS and the spleen, BM, mLN, and PPs were collected in cold PBS. Single-cell suspensions were prepared by mincing the spleens in PBS and filtered through 70 µm cell filters. Spleen samples were treated with erythrocytes lysis buffer and washed in PBS. The cells from PP were harvested by cutting the tissues with several strokes of sharp scissors and passing them through 30 µm cell strainers. Single-cell suspensions were kept on ice until further use. The total cell counts per organ were measured using an automated cell counter (Nexcelom Bioscience). From each sample, $5 \times 10^5$ cells were used to stain with fluorochrome-conjugated antibodies against cell surface markers, CD45 (30-F11), CD3 (17A2), CD19 (1D3), Ly6G (1A/8), IgG1 (RMG1-1), CD11b (M1/70), IgA (mA-6E1) and IgD (11-26C). All antibodies except CD11b (Thermo Scientific) were purchased from Biolegend (Germany). After washing, cells were suspended in flow cytometry buffer and measured on BD FACSAria$^{TM}$ flow cytometer and analyzed using FlowJo software (BD Biosciences).

**In vivo blockade of beta-adrenergic signaling and degradation of circulating DNA**

Mice were injected with non-specific beta-adrenergic receptor inhibitor Propranolol (Sigma; 30 mg/kg in 200 µl volume, i.p.). Compounds were diluted in sterile saline and administered 20 min after brain injury. For the degradation of circulating DNA, mice were injected with recombinant DNase-I (Cat. 11284932001, Roche; 1000 Units per mouse in 100 µl saline, i.v.) immediately after the onset of brain injury.

**Depletion of neutrophils and inhibition of NETs release**

Neutrophils depletion was achieved by the injection of anti-Ly6G antibody (Cat. BE0075-25, Bioxcell, 100 µg per mouse, i.p.) followed by anti-rat antibody (Cat. BE0122, 100 µg per mouse, i.p.) injection on the next day. Mice were once more injected with anti-Ly6g antibody (100 µg/mouse) before performing surgery. Mice were then sacrificed after 24 h for tissue analysis. The depletion of circulating neutrophils was verified by intracellular staining with fluorochrome-conjugated Ly6G (1A/8) antibody using FoxP3/transcription Factor staining kit (Thermo Scientific) followed by flow cytometry analysis. NETs release was inhibited via Cl-amidine, a pan PADs enzyme inhibitor. The stock solution of Cl-amidine (Cat. 506282, Millipore) was dissolved in dimethyl sulfoxide (DMSO, Sigma-Aldrich) and then further diluted in PBS. After brain injury induction, Cl-amidine was injected i.p. at 10 mg/kg.

**Optical clearing of Peyer's patches**

Mice were sacrificed and small intestines were dissected and placed in Petri dishes containing cold PBS. The intestinal tissue was then divided into four sections and cleaned from intestinal contents using cold PBS. About one cm of intestinal tissue containing PP was cut and further cleaned to remove food contents. Further, samples were embedded in 1.5% low melting agarose using plastic tissue molds, and samples were placed on ice. After 20 minutes, samples were carefully removed from the molds and extra solidified agarose was trimmed with a sharp surgical knife. Samples were then fixed with 4% PFA in PBS overnight at 4°C. The next day, samples were treated with a series of ethanol (EtOH) solutions starting from 30% and 60%, 80%, and 2 times 100%. Samples were kept in each solution for 12 h on an orbital shaker (50 rpm at 8°C) before a change to the higher percentage solution. At last, samples were treated with ethyl cinnamate (ECI, Cat. 112372-100G, Sigma) to match the refractive index and light-sheet fluorescence microscopy (LSFM) was performed.

**LSFM for PP volume analysis**

The cleared samples of PP were imaged using LSFM (UltraMicroscope BLAZE and UltraMicroscope II, Miltenyi Germany). The microscope systems employ the software ImSpector (LaVision BioTec, Germany). Samples were placed in the chamber filled with 100% ECI with the help of a steel holder. The autofluorescence of the sample was measured at 488 nm using an Optically Pumped Semiconductor Laser (OPSL; 50 mW) that was detected through a 525/50 nm bandpass filter. The light-sheet width has been set to 30% and the numerical aperture was set to 0.05. These settings correspond to a light-sheet thickness of 15 µm. The images were acquired with a zoom factor of 3.2X with an interval of 5 µm. For volume analysis, the images were processed using Imaris software version 9.5.1 (Bitplane, Switzerland). The Imaris File Converter was used to convert images. Software-own Gaussian and smoothing filters were used. The quantification of the volumes was performed with Imaris surface function and volumes were calculated. For this, every 25 µm area measurement of PP was performed in the autofluorescence channel.

**Whole-mount staining of PP and LSFM analysis**

Mice were sacrificed and small intestines were dissected and placed in Petri dishes containing cold PBS. Intestinal tissue proximal to the cecum was cut into three pieces; duodenum, jejunum, and ileum, respectively. Intestinal tissue, 5 mm proximal and distal to each PP, was cut and transferred to 4% PFA in separate 2 ml Eppendorf tubes and incubated for 2 h at RT. Samples were washed in PBS for 2 h and cleaned from intestinal contents and fat using PBS. Cleaned PP were transferred to 1 ml permeabilization buffer and incubated overnight on an orbital shaker (60 rpm). The following day, samples were transferred to a blocking buffer including 6% rat serum and incubated overnight on an orbital shaker (60 rpm) at 4°C. Further, samples were transferred into 2 ml Eppendorf tubes with blocking buffer containing 3% rat serum, and 1 µg purified anti-mouse CD16/32 antibody (BioLegend) for 30 minutes. Then, anti-CD19 Alexa Fluor 594 and anti-CD3 Alexa Fluor 647 (BioLegend) antibodies were added, and samples were incubated overnight at 4°C. Samples were transferred to new glass vials and washed with PBS for 1 h and then with freshly added PBS overnight with continuous shaking (60 rpm). The day after, samples were embedded in 1.5% low-gelling agarose using plastic tissue molds and put on ice for 20 minutes. Samples were then carefully removed from the molds and extra solidified agarose was trimmed with a sharp surgical knife. Samples in solidified agarose were transferred to fresh brown vials containing 2 ml 20% EtOH, followed by serial treatment of 40%, 60%, 80%, and 100% EtOH for 1 hour in each solution and then 100% EtOH incubation overnight. All steps were performed with continuous shaking of samples (60 rpm). Samples were finally transferred to ECI in fresh glass vials and imaged after 3 h.

The cleared and stained PP samples were imaged by LSFM (Ultramicroscope BLAZE, Miltenyi Biotech, Germany). Samples were placed in the chamber filled with 100% ECI solution with the help of a steel holder. B cells stained with anti-CD19 Alexa Fluor 594 were measured on 595/20 nm OPSL (Optically Pumped Semiconductor Laser; 50 mW) and were detected by a 650/50 nm band pass filter. T cells stained with anti-CD3 Alexa Fluor 647 were measured at 630/30 nm using an OPSL and were detected by a 680/30 nm band pass filter. The light-sheet width has been set to 40% and the numerical aperture was set to 0.05. These settings correspond to a light-sheet thickness of 4 µm The images were acquired with a zoom factor of 6.4 X with an interval of 5 µm. 3D images were prepared using Z-stacks in the software Imaris version 9.5.1 (Bitplane, Switzerland). The Imaris File Converter was used to convert the images.

**Analysis of stroke patient plasma**

The ethical approval for the use of healthy and stroke patients' plasma was granted as per the institutional ethics board committee of the University Hospital Essen (Study number: 18-8408-BO). Blood samples from stroke patients were taken at the University Hospital Essen Stroke Unit within three days after symptom onset. Samples were centrifuged at 900 g for 20 min, followed by a second centrifuge at 2000 g for 15 min to separate plasma. Aliquots from plasma were stored at –80°C until further analysis.

**Measurement of mouse plasma IgA, IgG, IgM and epinephrine**

EDTA blood was centrifuged at 8000 g for 10 min and plasma was collected in sterile tubes and frozen at –80°C until further use. Plasma samples were thawed and used to measure immunoglobulin levels using immunoassay kits as described by the manufacturer respectively, IgA (Cat. 88-50450-22, Thermo Scientific), IgG (Cat. 88-50400-22, Thermo Scientific) and IgM (Cat. 88-50470-22, Thermo Scientific). Plasma epinephrine was quantified using an immunoassay kit as described by the manufacturer (Cat. KA1877, Abnova).

**Quantification of plasma DNA**

The EDTA blood was centrifuged at 8000 g for 10 minutes and plasma was collected in sterile tubes and frozen at –80°C until further use. Samples are thawed and prepared with Qubit™ dsDNA HS– Assay-Kit (Cat. Q32851, Thermo Fisher Scientific). For total DNA quantification, samples were measured on Qubit Flex Fluorometer (Thermo Fisher Scientific).

**Quantification of NETs in plasma**

NETs quantification was performed on EDTA plasma using a previously described capture ELISA, which is based on citrullinated histone H3 associated with DNA (Sun et al., 2021). Anti-histone H3 antibody (5 µg/ml; ab5103, Abcam) was coated overnight at 4°C onto 96-well plates followed by 5% BSA blocking for 2 h. Wells were three times washed with 300 µl washing buffer followed by the addition of 50 µl plasma and 80 µl incubation buffer (including peroxidase-labeled anti-DNA antibody) for 2 hours at 300 rpm (Cell Death ELISA$^{PLUS}$, Cat. 11774425001, Roche). Then, the wells were washed three times with 300 µl washing buffer and 100 µl peroxidase substrate was added to the wells for 30 min in the dark. Afterward, 100 µl ABTS peroxidase stop solution was added to the wells, and absorbance was measured at 405 nm and was subtracted by absorbance at 490 nm (Abs 405 nm– Abs 490 nm). The absorbance values were considered in direct proportion to the amounts of soluble NETs and were presented as a relative increase to control.

**Isolation of B cells from PP**

Single-cell suspensions from PP were prepared and used for B-cell isolation using magnetic associated cell sorting (MACS). Briefly, cells were counted and resuspended in 35 µl MACS buffer. Afterward, B cells were purified using Pan B Cell Isolation Kit II, mouse based on the manufacturer’s instructions (Cat. 130-095-813, Miltenyi Biotec). This procedure provides B cells with very small amounts of unwanted intestinal epithelial and endothelial cells. To further enrich B cell preparations, cell suspensions were further purified using direct B cell labeling with CD45 microbeads using MojoSort™ Mouse CD45 Nanobeads (Cat. 480028, Biolegend). This isolation procedure provides B cells with a purity of ≥99%.

**RNA isolation and Illumina sequencing**

RNA was isolated from MACS-purified B cells using RNeasy Micro Kit according to the protocols provided by the supplier (Cat. 74004, Qiagen). Briefly, cells were lysed in lysis buffer on ice and centrifuged to remove cell debris. RNA was precipitated with 70% ethanol and passed through the RNA columns. After two washing steps, genomic DNA was removed using RNase-free DNase set (Cat. 9254, Qiagen). The columns containing RNA were washed with buffers and supplemented with centrifugation steps. In the end, RNA from columns was collected using RNase-free water in new Eppendorf tubes and stored at –80°C until further used for the analysis.

**RNA sequencing analysis**

The quality of sequenced reads was determined with FastQC (version v0.11.9) (Andrews, 2010) and Illumina adapters and low-quality reads -Phred score under 20- were trimmed with Trimmomatic (version 0.39) (Bolger et al., 2014). Kallisto (version 0.48.0-1) (Andrews, 2010) was used to pseudo-align reads to the GRCm38 release 102 genome assembly from ENSEMBL (Howe et al., 2021) and subsequently to quantify transcript expression as transcripts per million (TPM). Transcripts with less than 1 TPM across all 12 samples were removed and in R (version 4.1.2) (Team, 2021), a Wald Test from package Sleuth (version 0.30.0) (Pimentel et al., 2017) was used to analyze differential expression between the sham-operated and brain injured sample groups at the gene level, with significance set at a p-value < 0.05. Gene Ontology (GO) (Ashburner et al., 2000; Gene Ontology, 2021). Biological Process enrichment analysis was also performed in R, with package clusterprofiler (version 4.0.5) (Wu et al., 2021; Yu et al., 2012) and significance was defined as FDR-adjusted p-value < 0.05. In gene set enrichment analysis (GSEA), the dot size indicates the ratio of the overlap of differentially expressed genes with single GO term gene sets to the overlap of differentially expressed genes with the sum of all unique GO term genes queried.

**Cryo-sectioning of Peyer's Patches**

Murine ileum sections containing Peyer's Patches were isolated and fixed in 4% PFA/PBS (pH 7.4) for 2 h at 4°C. Tissue was then transferred to 15% sucrose/PBS (pH 7.4) and incubated at 4°C overnight, followed by a transfer to 30% sucrose/PBS (pH 7.4) and further incubation at 4°C overnight. Samples were embedded in Tissue-Tek O.C.T. Compound (Sakura Finetek) and 10 µm-thick histological sections were subsequently prepared on a CryoStar NX70 cryostat (Thermo Scientific) using Cryofilm Type 2C(9) (C-MK001-A2, Section-Laboratory).

**Histological immunofluorescence staining**

Cryo-sections were permeabilized and blocked with 1% BSA, 0.1% Tween-20, 0.1% DMSO/PBS (pH 7.4) for 1 h at room temperature. Two staining panels for the detection of either Ly-6G or cit-H3 were prepared. For Ly-6G staining, antibodies CD3-AF647 (17A2; 1:200), CD19-AF594 (6D5; 1:200), and Ly-6G-AF488 (1A8; 1:200; BioLegend) were diluted in blocking buffer and sections incubated with the antibody cocktail at 4°C overnight. For cit-H3 staining, antibodies CD3-AF647 (17A2; 1:200), CD19-AF594 (6D5; 1:200; BioLegend) and anti-cit-H3 (1:100; Abcam, ab5103) were diluted in blocking buffer and sections incubated with the antibody cocktail at 4°C overnight. After washing sections thrice with DPBS, donkey anti-rabbit antibody conjugated to Alexa Fluor Plus 488 (1:200; Invitrogen, A32790) diluted in blocking buffer was added and sections incubated for 2 h at room temperature. All sections were then washed thrice with DPBS, cell nuclei stained with DAPI (1:500; Carl Roth) for 10 min at room temperature in blocking buffer and washed again twice in DPBS. Sections were mounted in Dako Fluorescence Mounting Medium (Agilent Technologies) and dried for 4 h at room temperature in the dark.

**Confocal laser scanning microscopy of histological samples**

For high-resolution imaging of fluorescence-labeled Peyer's Patch samples, a Leica TCS SP8 confocal laser-scanning microscope with acousto-optic tuneable filters, an acousto-optical beam splitter, internal hybrid detectors (HyD SP), and a LMT200 high precision scanning stage was used. Overview images were constructed by acquisition of individual tiles using a Leica HC PL FLUOTAR 10x/0.30 DRY objective combined with a digital zoom factor of 1.2 and a resolution of 1024 x 1024 pixel per tile via the Leica Navigator module of the acquisition software (Leica LAS X). At the end of the tile acquisition, tiles were stitched automatically to create the overview image. Detail images were acquired via a Leica HC PL APO CS2 63x/1.30 GLYC objective combined with a digital zoom factor of 0.75 and a resolution satisfying the Nyquist criterion. In both cases, fluorescent signals were acquired using sequential scan mode. Alexa Fluor 488 (Plus) was excited via an argon laser at 488 nm and detected with an interal HyD at 500-550 nm. Alexa Fluor 594 was excited by a diode-pumped solid-state laser at 561 nm and detected with an internal HyD at 600-650 nm. Alexa Fluor 647 was excited with an helium-neon laser at 633 nm and detected with an internal HyD at 645-700 nm. Lastly, DAPI was excited by a diode-pumped solid-state laser at 405 nm and detected by an internal HyD at 450-500 nm. Detail images were subjected to deconvolution via Huygens Professional software (SVI) at default settings and reconstructed with Imaris software (Bitplane).

**Proteomics analysis of circulating neutrophils after brain injury**

Blood neutrophils were purified from sham-operated and brain-injured mice after 6 h using a neutrophil isolation kit (Miltenyi Biotech). Extracted protein from neutrophils was carbamidomethylated with 20 mM iodoacetamide (IAA) and digested with trypsin in 1:20 ratio using S-trap (micro column) digestion as described in the manufacturer's protocol. Briefly, proteins were acidified with 2.5% phosphoric acid. Acidified proteins were trapped onto the s-trap microcolumn with subsequent removal of contaminants using the S-trap binding buffer (90% MeOH and 10% TEAB, pH-7.6). Digestion of trapped protein was performed on column with trypsin followed by peptide elution with 50mM ABC, 0.1% formic acid, and 50% acetonitrile. The eluted peptides were further vacuum dried, and dissolved in 0.1% trifluroacetic acid for label-free quantitative proteomic analysis on Orbitrap Eclipse Tribid Mass spectrometer (Thermo Fisher Scientific) coupled to a nanoflow liquid chromatography system. After preconcentration of peptides on a trap column (C-18, 100 µm diameter and 2cm length, Acclaim Pepemap100) for 5 min, they were subsequently separated on C-18 reversed-phase nanocolumn with 75 µm diameter and 50-cm length (Acclaim PepMap100, Thermo Scientific). A gradient of 3–35% solvent B (B is 84% acetonitrile, 0.1% formic acid) over 90minutes was used for the peptide separation with a flow rate of ~400 nl $min^{-1}$. The eluted peptides from C-18 nanocolumn were subjected to nanospray ionization and analyzed by an orbitrap mass analyzer at a resolution of 12,000. The tandem mass spectra (MS/MS) generated after high-

energy collision dissociation of intense ions were analyzed along the chromatographic run by ion trap. All data were acquired in a data-dependent manner with 3 sec cycle time.
All mass spectrometric raw datasets were analyzed using Proteome Discoverer v.2.5.0.400 (Thermo Fisher). Briefly, MS/MS spectra were analyzed and quantified using an in-built search engine MASCOT (Thermo Fisher) against the UniProt mouse database (UP000000589, downloaded on 10.08.2019). The mass tolerance for parent ion and fragment ion was set at 10 ppm and 0.5Da respectively. The carbamidomethylation of cysteine was specified as static modification whereas oxidation of methionine was a dynamic modification. Master proteins having a maximum of 1% FDR and greater than or equal to 2 unique peptides were selected to increase data reliability. Missing values were imputed using low sampling abundance. The data was further normalized based on the total peptide amount and scaled before statistical analysis (i.e., t-test and fold change). The code for PCA plot is uploaded in GitHub (https://github.com/Susmita-isas/Stroke-mice-/new/main). The volcano plot was made using ggVolcanoR (https://ggvolcanor.erc.monash.edu/). Functional enrichment of statistically significant proteins was performed in g:Profiler (https://biit.cs.ut.ee/gprofiler/gost).

**Deep learning-based pipeline for B cell volume analysis of 3D images**

The volume of B cells and T cells was measured from the segmentation results of the corresponding channels of the 3D LSFM images. A human-in-the-loop deep learning-based segmentation method was employed to obtain the final segmentation results. The overall segmentation workflow was generalized from the iterative deep learning workflow in the Allen Cell and Structure Segmenter and demonstrated in Supplementary Fig. 1G. During this process, human experts were only asked to manually annotate two image z-stacks for the initial training of the deep neural network models, and then only to perform curation and minor error correction, to minimize the human effort. After the initial training, the model was applied to all images to generate segmentations, including post-processing steps, such as binarization and removing small false-positive objects from the model prediction. The results were provided to human experts to review. Segmentations either with negligible/minor errors or with big errors but straightforward to fix were selected. After fixing the errors, the curated subset was used to further train the segmentation models, which would then be applied to all images again. Such a loop will terminate when all segmentations were of good quality. Two different models were developed for B cells and T cells, respectively.

It is worth mentioning that there is no absolute per-pixel ground truth for the segmentation due to the limited resolution and diffraction of light. In other words, human experts can only tell if the segmentation is reasonable or not, but cannot be certain on every individual pixel if it belongs to B cells /T cells or not. For this reason, human curation (i.e. deciding if the preliminary segmentation is reasonable or not and making very minor corrections) is more useful and more efficient than human manual annotation of the images. It has been observed that the segmentation generated from the deep learning model was of better quality than human's direct annotation, especially in terms of 3D spatial consistency of the segmentation (e.g, a manually slice-by-slice annotated 3D ball-shaped object looks zig-zag from XZ or YZ side view, while the model's prediction is more consistent with a 3D ball-shaped). The models were able to learn useful knowledge from the noisy training data.

The images were down-sampled in XY dimensions by a ratio of 0.25. The neural network architecture was the enhanced 3D UNet, trained with a weighted sum of DICE loss and cross-entropy loss on a single NVIDIA A100 GPU. All details for model training and inference can be found at https://github.com/MMV-Lab/peyers_patch. The trained models are available at https://zenodo.org/record/6302990#.YhyocqvMI2x for reproducibility. For the final volume analysis, volumes in pixels obtained from volume analysis were converted to micrometer cubes, by multiplying with the voxel size of 0.975*0.975 at 5 μm intervals.

**Statistical analysis.** Data were analyzed using GraphPad Prism version 9.0. All datasets were tested for normality using the Shapiro–Wilk normality test. For comparisons between more than two groups, ordinary one-way ANOVA with Bonferroni's multiple comparison post-hoc tests was used. The comparisons between the two groups were analyzed using the two-tailed Mann–Whitney *U* test. Differences with *p*-values ≤ 0.05 were considered to be statistically significant.
