## Supplementary material for "Peyer’s Patch B cells undergo cell death via neutrophil-released toxic DNA following sterile tissue injury": Table S5

Supplementary table 5a. General neuroscore

| | | Time-point of scoring | score |
|---|---|---|---|
| **General Neuroscore** | **Hair** | 0. Hair neat and clean | |
| | | 1. Localized piloerection and dirty hair in 2 body parts (nose and eyes) | |
| | | 2. Piloerection and dirty hair in >2body parts | |
| | **Ears** (mouse on an open bench top) | 0. Normal (ears are stretched laterally and behind, they react by straightening up following noise) | |
| | | 1. Stretched laterally but not behind (one or both), they react to noise | |
| | | 2. Same as 1. NO Reaction to noise. | |
| | **Eyes** (mouse on OBT) | 0. Open, clean and quickly follow the surrounding environment | |
| | | 1. Open and characterized by aqueous mucus. Slowly follow the surrounding environment | |
| | | 2. Open and characterized by dark mucus | |
| | | 3. Ellipsoidal shaped and characterized by dark mucus | |
| | | 4. Closed | |
| | **Posture** (place the mouse on the palm and swing gently) | 0. The mouse stands in the upright position with the back parallel to the palm. During swing, it stands rapidly. | |
| | | 1. The mouse stands humpbacked. During the swing, it flattens the body to gain stability. | |
| | | 2. The head or part of the trunk lies on the palm | |
| | | 3. The mouse lies on one side, barely able to recover the upright position. | |
| | | 4. The mouse lies in a prone position, not able to recover the upright position. | |
| | **Total score for general scoring (normal=0 max=12)** | | |

Supplementary table 5b. Focal neuroscore

| Time-point of scoring | |
|---|---|
| **Body symmetry** (mouse on OBT, observe the nose-tail line) | 0. Normal (Body: normal posture, trunk elevated from the bench, with fore and hindlimbs leaning beneath the body. Tail: straight) |
| | 1. Slight asymmetry (Body: leans on one side with fore and hindlimbs leaning beneath the body. Tail: slightly bent.) |
| | 2. Moderate asymmetry (Body: leans on one side with fore and hindlimbs stretched out. Tail: slightly bent). |
| | 3. Prominent asymmetry (Body: bent, on one side lies on the OBT. Tail: bent) |
| | 4. Extreme asymmetry (Body: highly bent, on one side constantly lies on the OBT. Tail: highly bent) |
| **Gait** (mouse on OBT. Observed undisturbed) | 0. Normal (gait is flexible, symmetric and quick) |
| | 1. Stiff, inflexible (humpbacked walk, slower than normal mouse) |
| | 2. Limping, with asymmetric movements |
| | 3. Trembling, drifting, falling |
| | 4. Does not walk spontaneously (when stimulated by gently pushing the mouse walks no longer than 3 steps) |
| **Climbing** (mouse on a 45º surface. Place the mouse in the center of the | 0. Normal (mouse climbs quickly) |
| | 1. Climbs with strain, limb weakness present. |
| | 2. Holds onto slope, does not slip or climb |
| | 3. Slides down slope, unsuccessful effort to prevent fail |
| | 4. Slides immediately, no effort to prevent fail. |
| **Circling behavior** (mouse on OBT, free observation) | 0. Absent circling behavior |
| | 1. Predominantly one-side turns. |
| | 2. Circles to one side, although not constantly. |
| | 3. Circles constantly to one side. |
| | 4. Pivoting, swaying, or no movement. |
| **Forelimb symmetry** (mouse suspended by tail) | 0. Normal |
| | 1. Light asymmetry: mild flexion of contralateral forelimb. |
| | 2. Marked asymmetry: marked flexion of contralateral limb, the body slightly bends on the ipsilateral side. |
| | 3. Prominent asymmetry: contralateral forelimb adheres to the trunk. |
| | 4. Slight asymmetry, no body/limb movement. |
| **Compulsory circling** (forelimbs on bench, hindlimbs suspended by the tail: it reveals the presence of the contralateral limb palsy) | 0. Absent. Normal extension of both forelimbs. |
| | 1. Tendency to turn to one side (the mouse extends both forelimbs, but starts to turn preferably to one side) |
| | 2. Circles to one side (the mouse turns towards one side with a slower movement compared to healthy mice) |
| | 3. Pivots to one side sluggishly (the mouse turns towards one side failing to perform a complete circle) |
| | 4. Does not advance (the front part of the trunk lies on the bench, slow and brief movements) |
| **Whisker response** (mouse on the OBT) | 0. Normal |
| | 1. Light asymmetry (the mouse withdraws slowly when stimulated on the contralateral side) |
| | 2. Prominent asymmetry (no response when stimulated to the contralateral side) |
| | 3. Absent response contralaterally, slow response when stimulated ipsilaterally. |
| | 4. Absent response bilaterally |
| **Total score for focal deficits (normal=0 max=28)** | |
